# A novel single layer microfluidic device for dynamic stimulation, culture and imaging of mammalian cells

**DOI:** 10.1101/2022.03.14.484262

**Authors:** Adil Mustafa, Elisa Pedone, Antonella La Regina, Ahmet Erten, Lucia Marucci

## Abstract

The possibility to tightly control the cellular microenvironment within microfluidic devices represents an important step forward precision analysis of cellular phenotypes *in vitro*. In particular, microfluidic platforms that allow both long-term mammalian cell culture and dynamic modulation of the culture environment can support quantitative studies of cells’ responses to drugs. Here, we report the design and testing of a novel microfluidic device of easy production (single Polydimethylsiloxane layer), which integrates a micromixer with vacuum-assisted cell loading for long-term mammalian cell culture and dynamic mixing of 4 different culture media. Finite element modelling was used to predict the flow rates and device dimensions to achieve diffusion-based fluid mixing. The device showed efficient mixing and dynamic exchange of media in the cell trapping chambers. This work represents the first attempt to integrate single layer microfluidic mixing devices with vacuum-assisted cell loading systems for mammalian cell culture and dynamic stimulation.

## Introduction

Culturing cells *in vitro* is of paramount importance in molecular biology to study cellular processes such as cell-cell interactions, responses to external stimuli (physical or chemical), acquisition of genetic mutations over time and so on^1–3^. Traditional *in vitro* methods for cell culturing (i.e., in dishes) do not allow spatio-temporal control of the cellular environment and are therefore limited when it is important to study phenotypic changes upon dynamic perturbations in the culturing conditions. Microfluidic devices find their advantage in using low sample volumes, while allowing precise control over the cell culture environment and mimicking of physiological conditions^4–7^. Recently, there has been an increasing interest to employ microfluidic devices for a range of mammalian cells applications, such as high throughput single cell studies^8–10^, parallelized drug screening^11^, temporal modulations of inputs^12^, easy-to-use perfusion for cell proliferation and differentiation studies^13,14^ and automated feedback control of gene expression^15^.

Dynamic drug stimulation, drug mixing and long-term media perfusion can be of great importance for cell biology studies. This is often achieved by integrating in microfluidic devices active and passive micromixers^16,17^, and using multiple layers (which come at the cost of sophisticated mold preparation) and/or separate devices for multiplexer input control and cell growth, respectively, as in^11^. Trenzinger and colleagues^18^ reported an automated microfluidic device with 48 independent chambers for both adherent and suspension cell cultures in rapidly changing media conditions; in this case, mixing was implemented on chip, but this device is still quite complex to fabricate as it includes valves and multiple layers.

Kolnik et al.^12^ designed a multilayer microfluidic device with a built-in function generator to create chaotic mixing between two different cell culture media. The device also implemented vacuum-assisted loading of cells in individual trapping chambers to protect them from the shear stress.

Single layer microfluidic platforms have been proposed to simplify fabrication and operation, and to facilitate cell loading^19,20^. Usually, such devices allow long-term mammalian cell culture but lack the ability of dynamically mixing two or more culture media.

In this study, we present a novel Polydimethylsiloxane (PDMS), single layer microfluidic device that can be employed to culture, for long term, mammalian cells while perfusing them with combinations of up to four cell culture media (possibly enriched with drugs). The device embeds a cell loading part, a diffusive mixing part and five cell culture chambers, adjoined to the perfusion channel (Figure 1). As in^12^, vacuum-assisted cell loading is employed. Media is provided using four inlets, each of them automatically regulated by software-controlled syringe pumps and connected to flow stabilizers and mixing serpentines structures. We show, both using finite element analysis and with experiments, that the device allows mixing of different inputs and dynamics change of the culture media provided to cells. Finally, the device allowed long-term culturing of two different mammalian cell lines. Our new device has specific advantages such as single layer fabrication, diffusion-based mixing, and easy cell loading. To our knowledge, this is the first study to use a single layer microfluidic device with an integrated vacuum-assisted cell loading mechanism. It should be possible to easily modify the device to increase its drug screening capability by modifying the number of inlets and serpentines.

**Figure 1.**
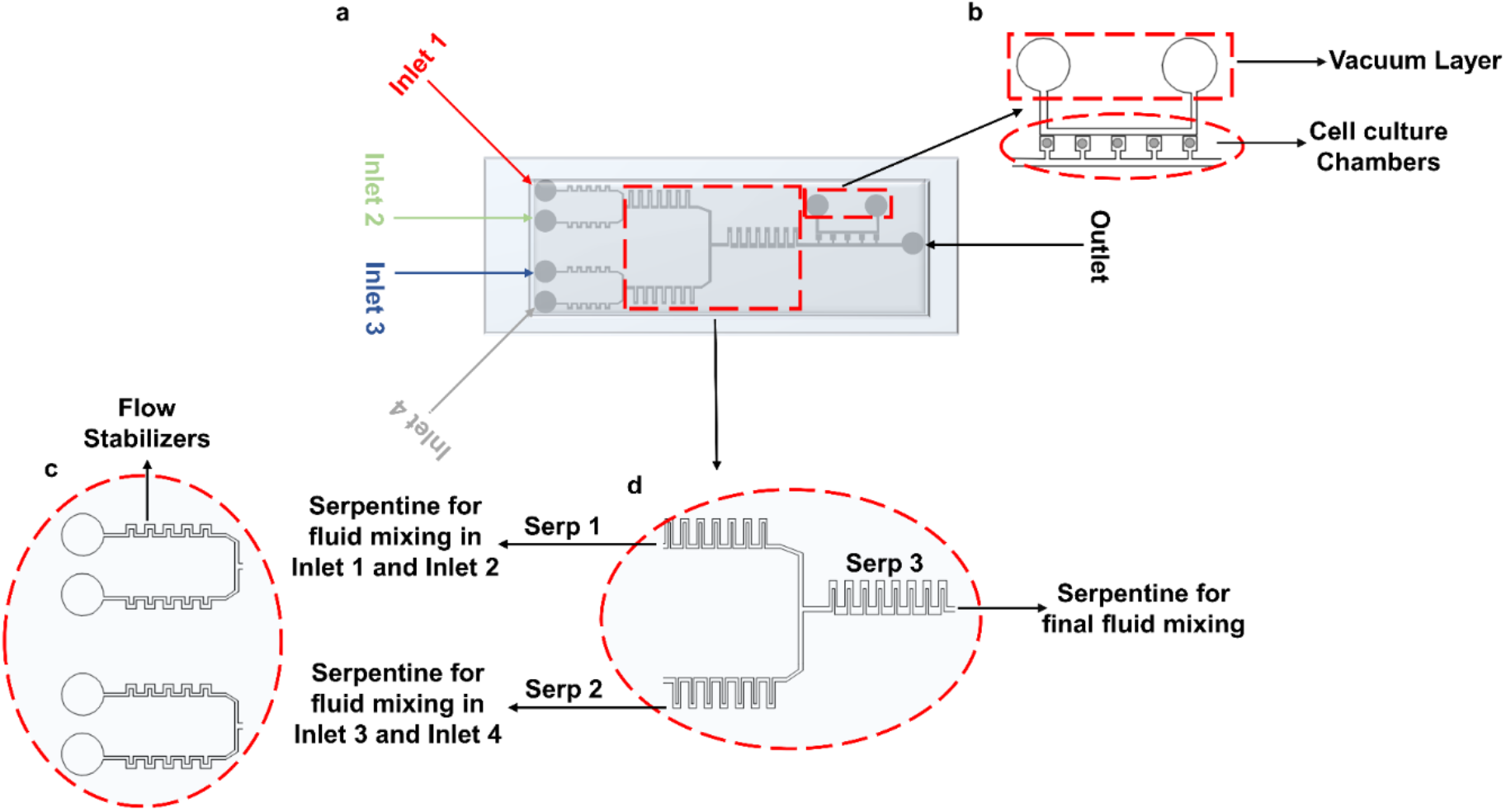
Microfluidic device design. (a) Graphical representation of the device bonded on glass slide. (b) Vacuum layer and cell culture chambers. (c) Inlet part of the device with flow stabilizing serpentines. (d) Mixing serpentines 1 and 2, which facilitate diffusive mixing of cell media and drugs; serpentine 3 enables diffusive mixing of culture media coming from serpentines 1 and 2.

## Materials and methods

### Device design

The device consists of 5 consecutive cell culture chambers (footprint 360 μm × 360 μm, Figure 1a). The ‘opening’ that connects the chambers to the main channel is 200 μm × 200 μm. The height of the channel is 50 μm. Cells are loaded using a vacuum pump connected to the vacuum layer running parallel to the main channel, as shown in Figure 1b. Figures 1 c and d represent the flow stabilizers and serpentine structures for fluid stabilization and mixing, respectively. The distance between the vacuum layer and the trapping chambers is 80 μm. A combination of different cell culture media/drugs can be infused through the device using four inlets, as shown in Figure 1. Inlets 1 and 2 come together in the top serpentine (Serp 1) structure, and Inlets 3 and 4 come together in the bottom serpentine (Serp 2) for mixing purposes. A larger serpentine structure (Serp 3) is connected at the junction of Serp 1 and Serp 2 for further mixing of the fluids coming from them. The dimensions of the serpentine, the number of the serpentines required to achieve a complete mixing, the fluid flow rates and the number of cell culture chambers were chosen given results from COMSOL simulations.

### Device fabrication

The microfluidic device reported in this study was fabricated using maskless photolithography. Computer aided design software (CAD) AutoCAD was used to design the input files; a negative photoresist SU-8 2050 (MicroChemicals GMBH, Stuttgart, Germany) was spin-coated on a 4-inch silicon wafer (Figure 2a) at 3000 rpm for 30 seconds to achieve a height of 50 μm, as shown in Figure 2b. A pre-exposure bake was performed in two steps by placing the wafer directly on to the hot plate at 65 °C for 2 minutes and then at 95 °C for 9 minutes.

**Figure 2.**
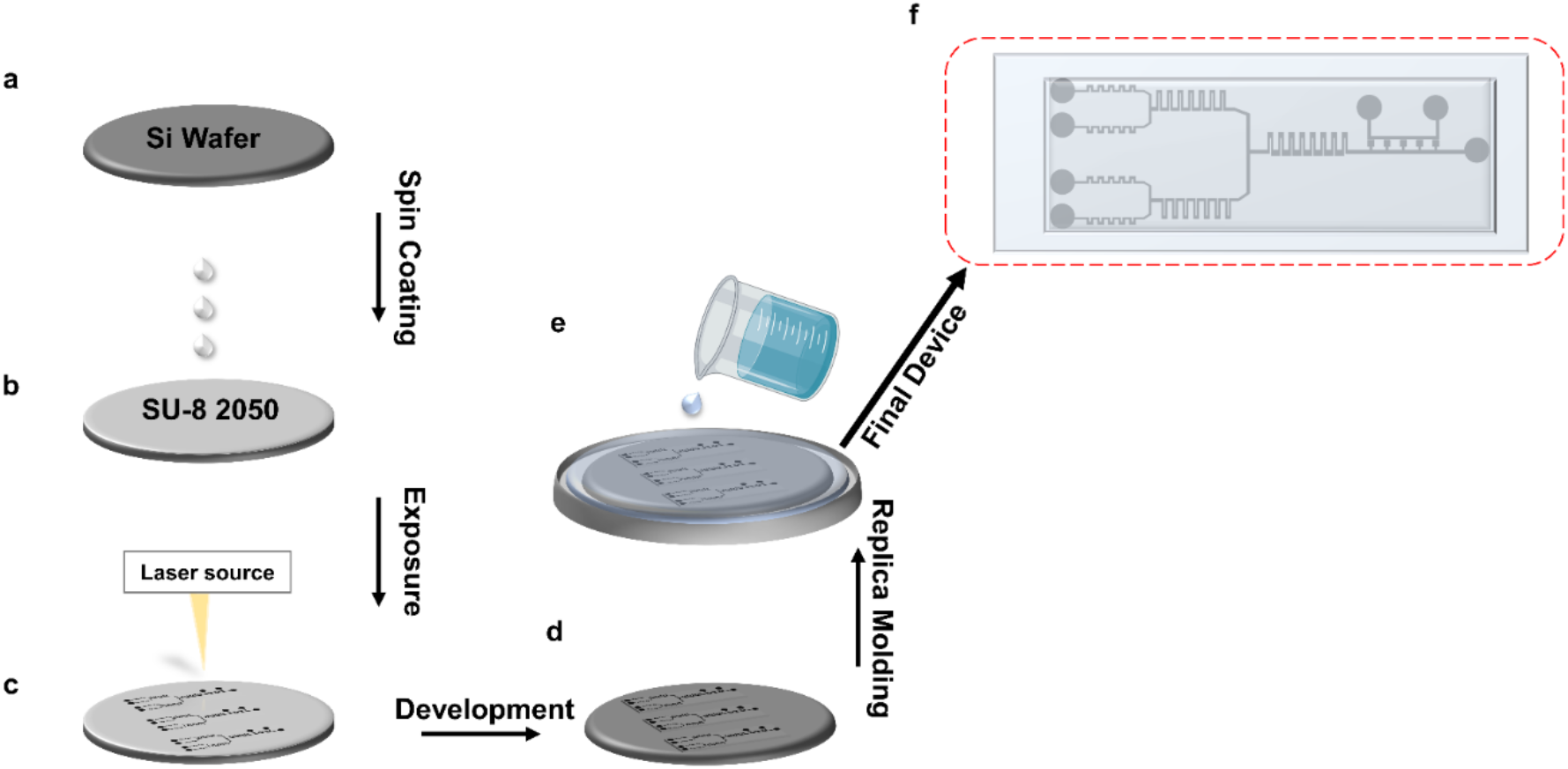
Device fabrication protocol. (a) 4-inch silicon wafer. (b) Spin coating SU-8 2050 negative photoresist on 4-inch silicon wafer. (c) Exposure of spin-coated silicon wafer using laser source. (d) Removal of excess negative photoresist using SU-8 developer solution. (e) PDMS replica molding. (f) Final fabricated device bonded on glass slide.

The baked silicon wafer was then exposed using a Heidelberg laser writer ‘μPG101’ (Heidelberg Instruments Mikrotechnik GmbH) using 30 mW of power at 10 % (Figure 2c). A post-exposure baking step was performed first at 65 °C for 1 minute and then at 95 °C for 7 minutes. The exposed wafer was than immersed in the SU-8 developer solution (MicroChemicals GMBH, Stuttgart, Germany) to remove the access photoresist. The structure was obtained in the development step (Figure 2d), and we refer to it as the ‘mother mold’. To further enhance the adhesion of structures to the wafer, a hard baking was done by placing the mother mold directly on to the hot plate at 200 °C for 10 minutes.

The final device was obtained by pouring PDMS on the mother mold with a base to curing agent ratio of 10:1 (Figure 2e). A degassing step was performed to remove the air bubbles by placing the mold with PDMS in a desiccator for 25 minutes. After degassing, the mother mold was placed in an oven at 70 °C for 120 minutes. The baked PDMS mold was autoclaved to achieve sterilization. The individual chips were realized by bonding the PDMS devices on a microscope glass slide (75 mm × 25 mm) using oxygen plasma at 100 % power as shown in Figure 2f.

### Cell culture and device loading

Two mouse Embryonic Stem Cell (mESC) lines (REX-dGFP2^21^ and MiR-290-mCherry/miR-302-eGFP^22^) were used to test cells’ viability in the microfluidic device. Before loading in the chip, mESCs were cultured on gelatinized tissue culture dishes at 37 °C in a 5% CO2 humidified incubator in Dulbecco’s modified Eagle’s medium (DMEM) supplemented with 15% fetal bovine serum (Sigma), non-essential amino acids, L-glutamine, sodium pyruvate, Penicillin-Streptomycin, 2-mercaptoethanol and 10 ng/ml mLIF (Peprotech; 250-02). Once the mESCs in culture dishes reached confluency, they were detached using 3 mL Gibco trypsin-EDTA (0.5%). 1 mL of detached cells suspended in trypsin were collected in a falcon and centrifuged at 1200 RPM for 5 minutes. The supernatant was removed using a micropipette and the cells were then resuspended in 100 μL of the cell culture media. The cells were then loaded into a 2.5 mL syringe avoiding any bubbles. The cell loading into the microfluidic device was done in two steps. First, the chip was wetted by using the cell culture media. In the next step, the syringe with cells was attached to the outlet and cells were infused into the chip. Once the channel was filled with the cells, a vacuum pump at a pressure of −80 kPa connected to vacuum layer was switched on to aspirate the cells into the chambers and to remove any air in the chambers. Once loaded, the device was infused with fresh media to clear the excess cells.

### Mixing experiments and imaging

For experiments in Figures 4-7 and in Figure S2, media was enriched with the fluorescence dyes Atto 488 (41051-1MG-F), Atto 647 (97875-1MG-F), Rhodamine B CAS-81-88-9 and Phosphate-buffer saline (D8537-500 ML), all obtained from Sigma Aldrich, and used at a 10uM concentration. For media delivery, we used syringe pumps (World Precision Instruments AL-1000) as shown in Figure 3. A customized Matlab code was used to run the pumps and change the flow rates during dynamic mixing and media exchange experiments.

**Figure 3.**
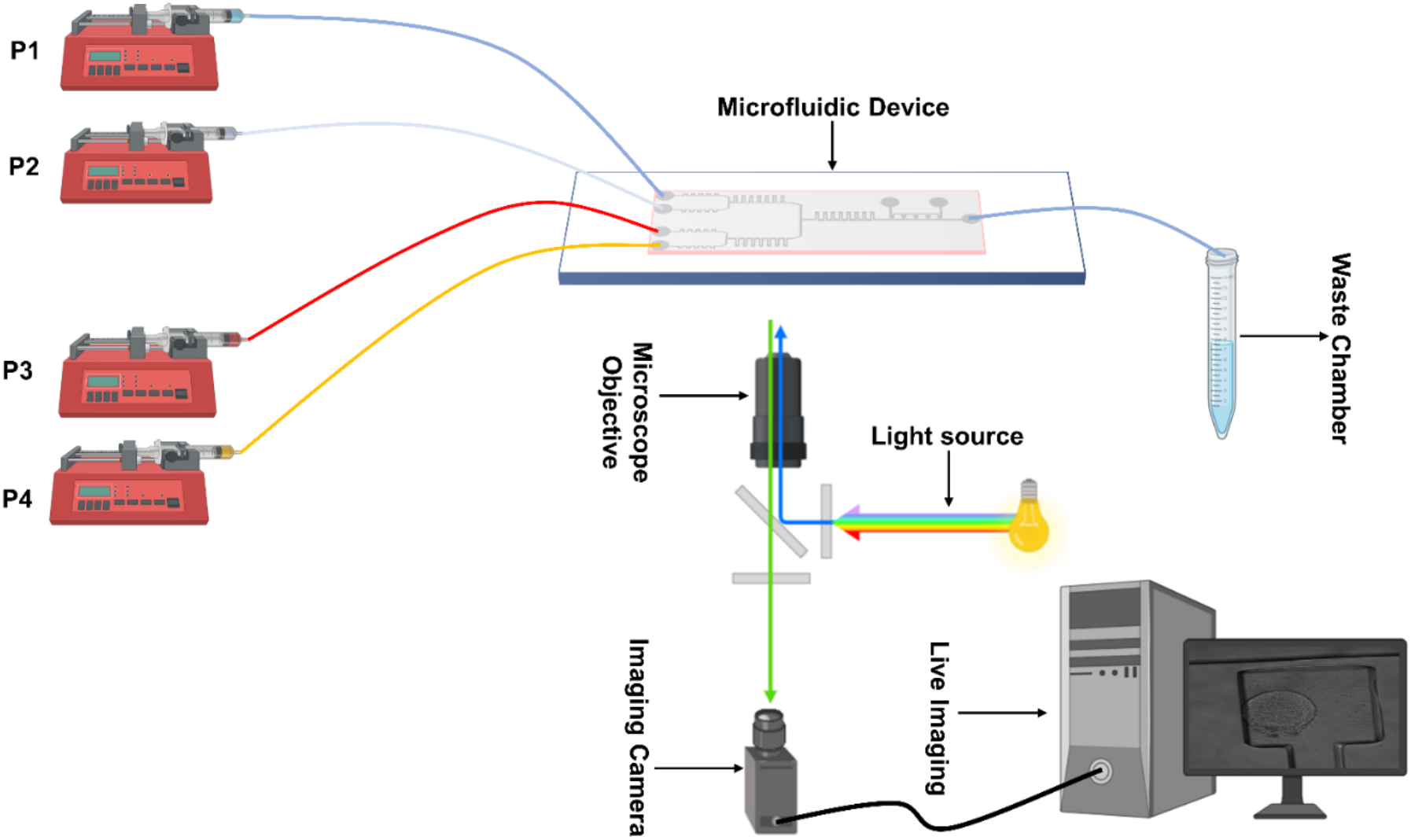
A schematic of the experimental setup for cell culturing and imaging on-chip.

**Figure 4.**
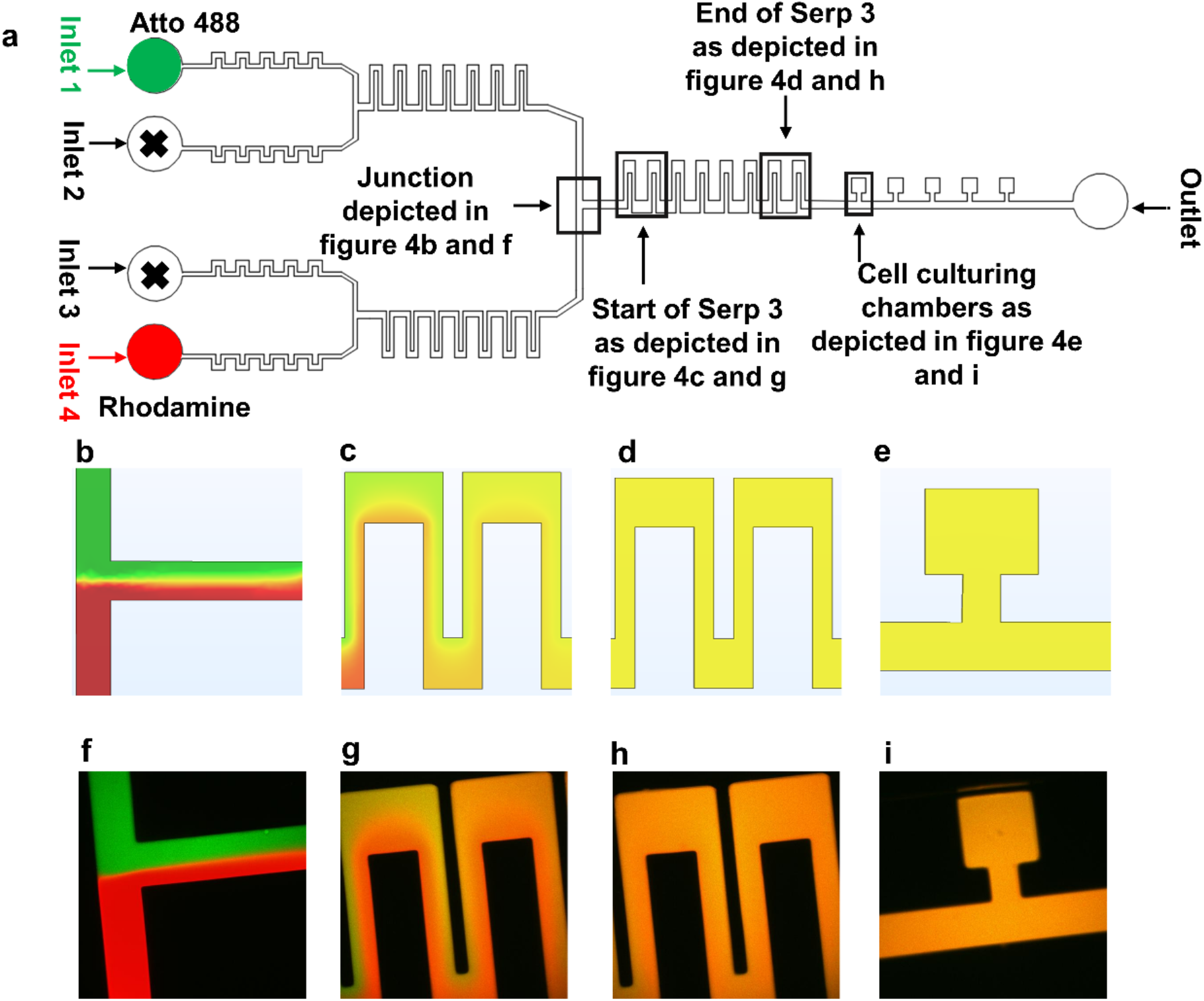
Mixing 2 different fluids. The flow rate used for these experiments was 50 μl hr^-1^. (a) Microfluidic device with highlighted parts. (b-e) COMSOL simulation depicting diffusion-based mixing of two fluids using inlets 1 and 4. (f-i) Experimental results for static mixing using Atto 488 and Rhodamine B dyes at 50 μl hr^-1^.

The imaging experimental setup consisted of Leica LASX live cell imaging workstation on DMi8 inverted fluorescence microscope (Leica Microsystems Wetzlar, Germany). An Andor iXON 897 Ultra back integrated camera was used for imaging. Images collected to test the efficacy of the device for the media exchange and mixing were recorded by using an objective lens of magnification 10x; cell viability experiments’ images were collected using a 20x magnification. The media exchange and mixing experiments in Figures 4-6 were 30 minutes long with images recorded every 2 minutes, while the experiments in Figure S2 were 145 minutes long with images recorded every 5 minutes. For cell viability experiments, microscopy images were collected every 24 hours.

The experiments where the fluid concentrations were changed dynamically required balancing the flow rates in the main junction as, in the device, the total flow rate at inlet 1 and 2 should always be equal to the total flow rate at inlets 3 and 4. Such balancing avoids any back flow from inlets 3 and 4 towards the inlets 1 and 2. For example, if the flow rate at inlet 1 and inlet 2 is 50 μl hr^-1^ in each (100 μl hr^-1^ in total), then, in order to have no back flow, the total flow rate at inlets 3 and 4 should also be equal to 100 μl hr^-1^. Thus, in experiments where the flow rate at one of the inlets was changed dynamically, the flow rate in the other inlet(s) was adjusted (results from an exemplar experiment are provided in Figure 7).

The chip wetting process for the experiments where only two of the four inlets were used was handled carefully. Initially, wetting was done by keeping the inlets 2 and 3 open; the pumps connected to inlets 1 and 4 were started at high flow rates (i.e., between 6-10 ml hr^-1^) to fill the entire device. This resulted in fluids from inlets 1 and 4 flowing out of the open inlets 2 and 3. Afterwards, these inlets were sealed using knotted tubing. This procedure allows to run long-term cell culture experiments by minimizing air bubble generation.

### Diffusion and flow modeling

Fabrication of microfluidic devices requires resources and time not readily available. Hence, it is of utmost importance to that device design is optimized using all the available tools before the fabrication process is started. Computational fluid dynamics helps in the design optimization of the microfluidic devices. To verify the operating parameters of our device we used COMSOL 5.6 Multiphysics software. The device design generated by AutoCAD (Figure 4a) was imported into COMSOL 5.6 for the simulation purpose. The computational fluid dynamic (CFD) model was based on solving fluid flow by using laminar form of Navier-Stokes equation and modelling transport mechanism using mass transport equations. The fluid flow was modelled using ‘laminar flow’ built in Physics while transport mechanism was solved using the ‘transport of diluted species’ interface. Simulations were setup using fully developed flow boundary condition at the inlets to accurately model the laminar flow regime; the outlet boundary condition was set to be zero pressure. To validate the experiments, the dyes were used at the same concentration of the experiments in all simulations. The diffusion coefficient values for Rhodamine B and Atto 488 were used to simulate diffusive mass transport in the device. The governing equations, fluid velocity magnitude and shear stress in serpentine structures, connecting channel and cell culture chambers are provided in supporting information.

## Results and Discussion

### Diffusion Mixing Experiments

Firstly, we tested both via simulations and with experiments the device performance in mixing media from the different inlets (first just two, and then four). The finite element method was used to model the fluid flow and the mixing in the microfluidic device.

The simulation results revealed that the rectangular serpentine structures act as effective micromixers for two inputs, as shown in Figure 4a-e. We then tested experimentally the mixing of two media, containing Atto 488 (green dye) or Rhodamine B (red dye), infused with flow rates between 50-–100 μl hr^-1^ in inlets 1 and 4; inlets 2 and 3 were not used in these experiments and were closed (Figure 4a). It was observed that, at 50 μl hr^-1^, the two fluids flowed side by side in the initial part of the Serp 3 (simulation and experiment in Figure 3c and g, respectively) and mixed completely as they reached the end of Serp 3 and, ultimately, the cell culture chambers (simulation and experiment in Figure 4d, e, and h, i, respectively). Mixing at flow rates higher than 100 μl hr^-1^ might be possible by increasing the number of serpentine structures.

The device performance was further tested by mixing four different fluids; this was achieved by combining fluids in 2×2 configurations as shown in Figure 5a. The experiments were setup by keeping the flow rate at 50 μl hr^-1^ at all the four inlets. The fluids coming from inlet 1 (Atto 647, blue dye) and inlet 2 (phosphate-buffered silane, PBS) were combined in the top junction as shown in Figures 5b (simulation) and h (experiment), and they were mixed via serp 1. Similarly, fluids coming from inlet 3 (Rhodamine B, red dye) and inlet 4 (Atto 488, green dye) came together at the bottom junction (Figures 5c and i, simulation and experiment respectively) and were mixed via Serp 2. The final mixing step combined the mixed fluids at the junction of Serp 1 and Serp 2 using Serp 3.

**Figure 5.**
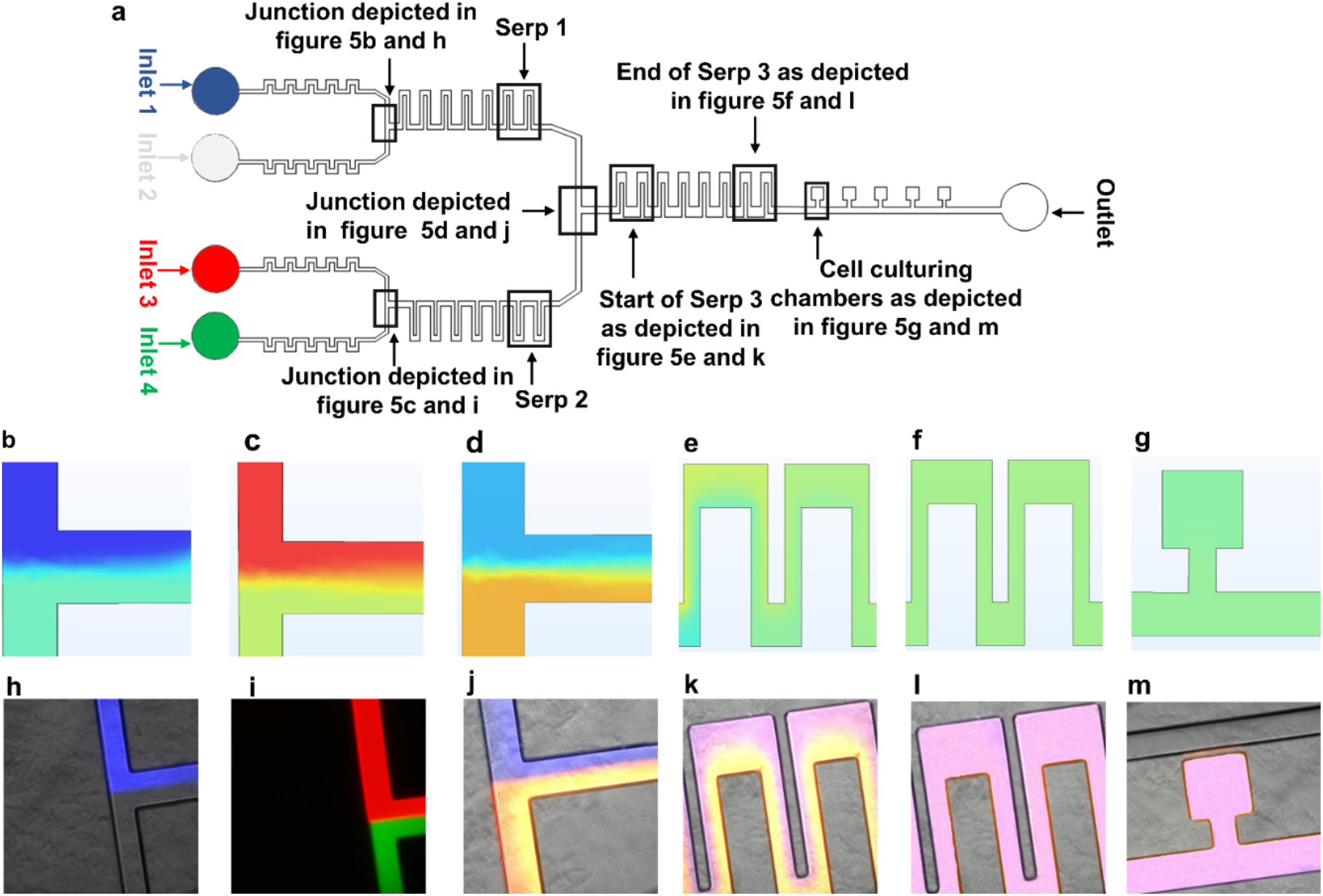
Mixing 4 different fluids. The experiments were designed to test the ability of the device to achieve mixing using serpentine structures. The flow rate at all the inlets was set at 50 μl hr^-1^. (a) Microfluidic device highlighting different parts. (b-g) COMSOL simulation depicting diffusion-based mixing of four fluids using inlets 1-4. (h-m) Experimental results for static mixing using Atto 488, Rhodamine B, Atto 647 and PBS at 50 μl hr^-1^.

As observed when mixing two fluids, there was a thorough mixing of the four fluids in Serp 3, and in the cell culture chambers (Figures 5 g and m, simulation and experiment, respectively). Differences in the mixed media color measured in experiments/simulations are because the colorless media used in experiments could not be represented in COMSOL, where a light blue color was instead used, and a light green color represented the Atto 488 dye.

We then tested our device for effective media exchange. We firstly set up simulations and experiments by using only two inlets, infused with two fluorescence dye solutions (Rhodamine B (red) and Atto 488 (green)) in PBS (Figure 6a). Initially, the pump controlling the syringe with the green dye was set at 50 μl hr^-1^ and the pump controlling the red dye-containing syringe was switched off. In this case, the whole microfluidic device, including cell culture chambers, was filled with the green dye (t=0 min, Figures 6b -simulation- and e -experiment-). We then (t=4 min) switched off the pump infusing the green dye and switched on the pump with the red dye at 50 μl hr^-1^. At t=6 min, the red dye started to take over the green one in the junction (Figures 6 c, f), taking over the perfusion channel and the cell culture chambers at t=8 min (Figures 6 d, g, j). Quantification of the fluorescence signals confirmed successful media exchange, with a short time delay between pumps’ switching and complete change of dye in the cell culture chambers (Figures 6 k and l). We also confirmed robust media exchange, mixing and flow stability in longer experiments (2 hours and 40 minutes, Supplementary information S2).

**Figure 6.**
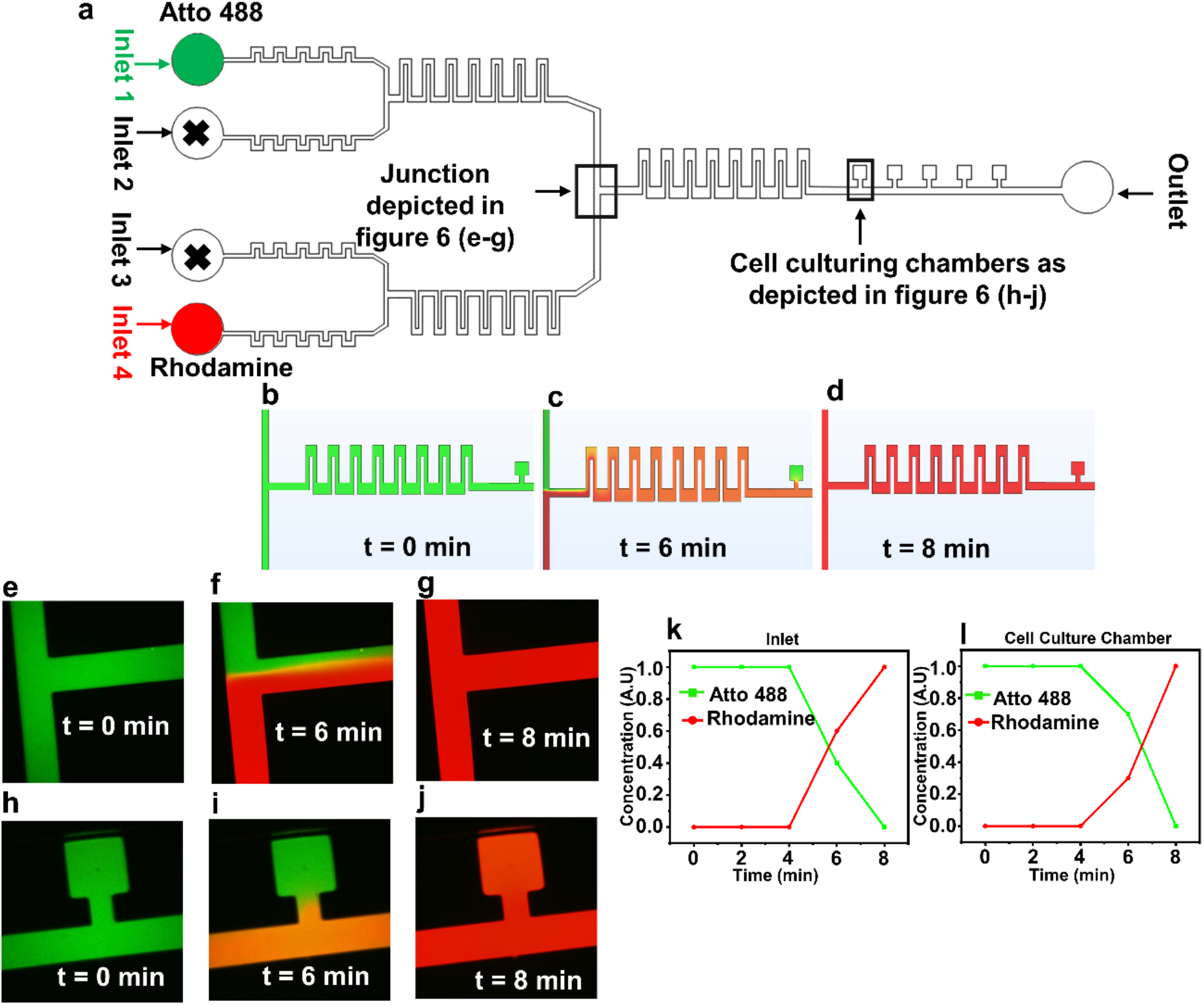
Media exchange experiments using Atto 488 and Rhodamine B. The experiments were designed to test the dynamic media exchange ability of the device by using two inlets. (a) Microfluidic device with highlighted parts. (b-d) COMSOL simulation depicting diffusion-based media exchange between two fluids using inlets 1 and 4. (e-j) Experimental results depicting media exchange in time. (k-l). The concentration plots were generated by quantifying the dye concentration at the inlet junctions and within the cell culture chambers.

The device performance was further assessed by setting up media exchange experiments using all 4 inlets (Figure 7). As in the two dyes case, COMSOL simulations (Figures 7 b-d) were performed. Inlets 1-4 were connected to the pumps infusing media containing Atto 647 (blue), PBS (colorless), Atto 488 (red) and Rhodamine B (green), respectively. Initially, all the four solutions were infused at 50 μl hr^-1^ (time t = 0 min, Figures 7b, e, h, k). The pump infusing the blue dye (inlet 1) was then completely switched off and the flow rate at the pump infusing PBS (inlet 2) was increased to 100 μl hr^-1^ to balance the flow rates at inlets 3 and 4. At t = 4 min, PBS started taking over the blue dye at the main junction (Figures 7c and f), with the color of the mixed fluid altered at the end of serp 3 and in the cell culture chamber (Figures 7i and l). Complete media exchange was obtained at t = 8 min (Figure 7d, g, j and m). Quantification of media fluorescence confirmed successful dynamic exchange of mixed fluid, with a short delay between the time of pump switching and the replacement of the media in the cell culture chamber.

**Figure 7.**
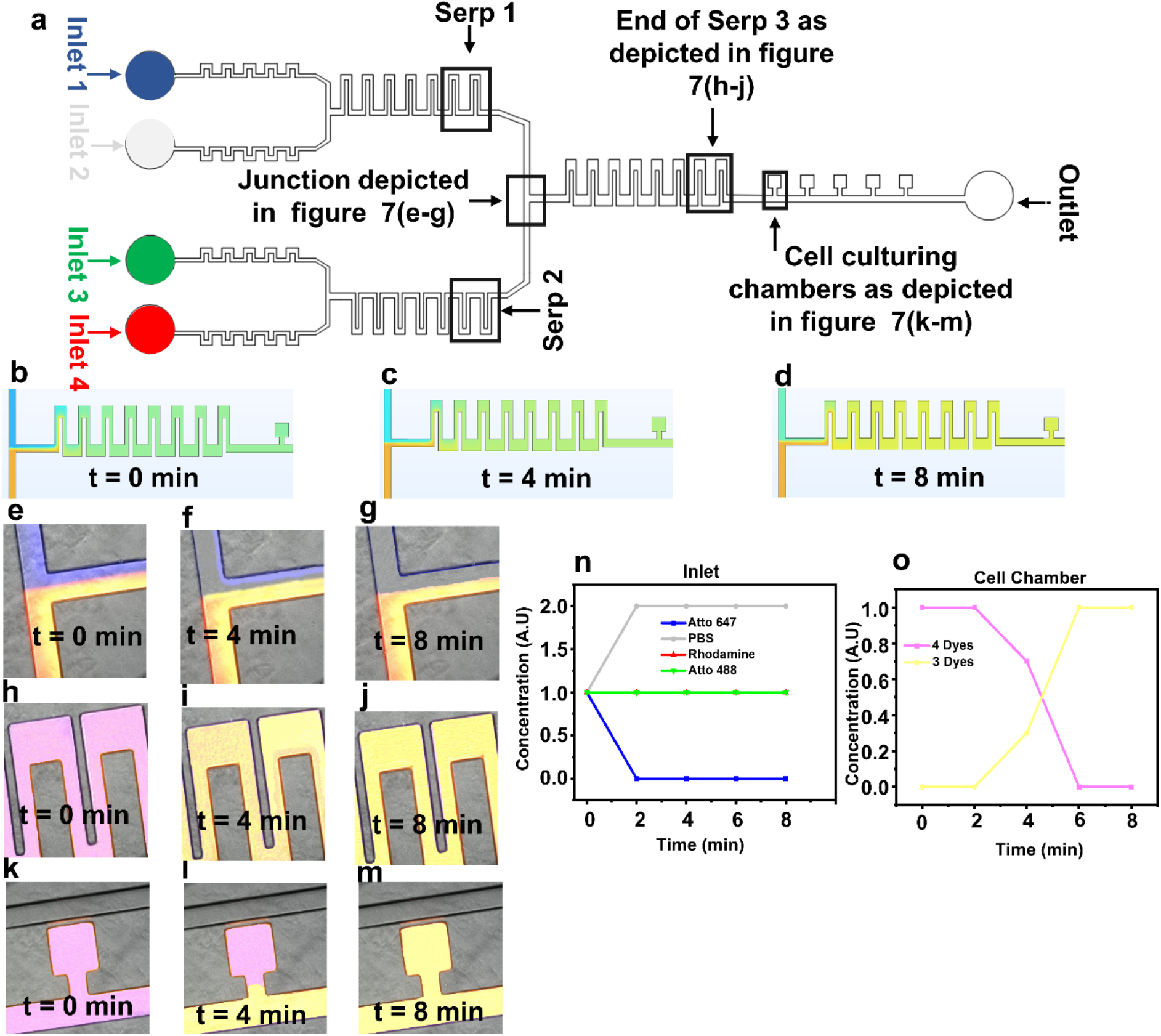
Media exchange experiments using Atto 647, PBS, Atto 448 and Rhodamine at 50 μl hr^-1^. The experiments were designed to test the dynamic media exchange ability of the device. (a) Microfluidic device with highlighted parts. (b-d) COMSOL simulation depicting diffusion-based media exchange of four fluids. (e-m) Experimental results depicting media exchange in time. The blue dye was switched off at t=0 min and the flow rate for PBS was increased to 100 μl hr^-1^. (n-o) Time vs concentration plots at the inlet junctions (n) and cell culture chambers (o). As the concentration of the green and the red dyes in the channel were never changed (fixed at 50 μl hr^-1^), they are represented by the concentration value of 1 (n). The concentration of PBS in the channel doubled as the flow rate was increased to 100 μl hr^-1^ for flow balancing purposes; this is represented by the concentration value of 2 (n).

### Cell Viability Experiments

We finally performed cell viability experiments to assess the suitability of our microfluidic device for long-term mammalian cell cultures. REX-dGFP2^21^ and MiR-290-mCherry/miR-302-eGFP (named here DRC)^22^ mouse embryonic stem cell lines were used for this purpose. The microfluidic devices loaded with cells were incubated at 37 °C and perfused with continuous media flow (syringe pump at 50 μl hr^-1^). The results in Figure 8 clearly show that both cell lines remain healthy and expand in our microfluidic device for at least 5 days.

**Figure 8.**
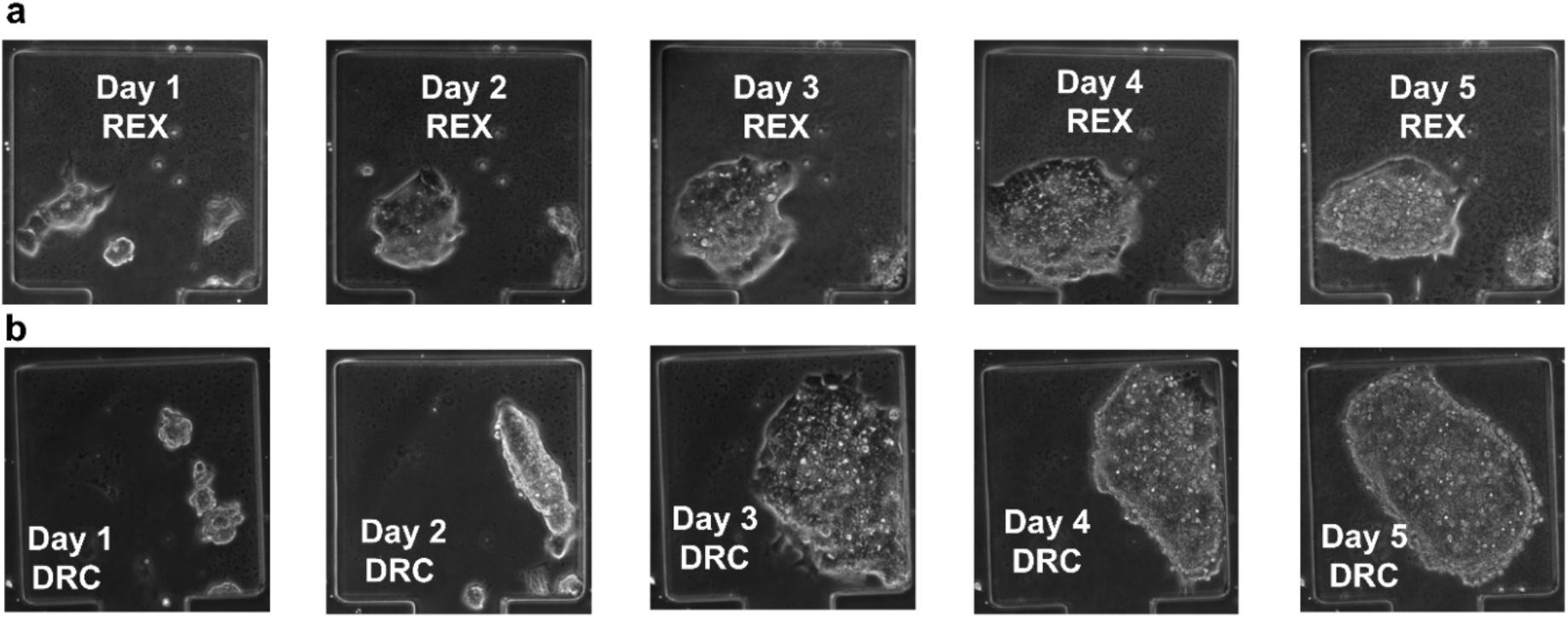
(a, b) Cell viability tests for REX-dGFP2 (REX, a) and MiR-290-mCherry/miR-302-eGFP (DRC, b) mouse embryonic stem cell lines.

## Conclusions

Exposing cultured cells to a specific media for a determined amount of time is of utmost importance for many biomedical and biological studies^23,24^. It is therefore necessary to have a controlled and rapid mechanism of media exchange for microfluidic devices that are designed and fabricated for such experiments. In this study, we presented a novel single layer microfluidic device that combines a passive microfluidic micromixing mechanism with vacuum-assisted cell loading for long-term mammalian cell culture. Our device allows cell culture with reduced shear stress and dynamical cell stimulation with combinations of 4 different media, which can be automatically delivered to cells via computer-controlled pumps. The mixing of the fluids is achieved by rectangular serpentine structures at fluid flow velocities in the μl hr^-1^ range. Finite element method simulations and experiments showed successful fluid mixing and media exchange in cell trapping chambers at flow velocities between 50 -100 μl hr^-1^, avoiding shear stressing of cells. Long-term cell culture (5 days) of two different mammalian cell lines, in a fully controllable culture environment, was achieved.

Our device, as compared to previously reported ones, is easier to fabricate and use, being single layer. Our design, (available at https://github.com/LM-group/Single-layer-microfluidic-device-), could be modified for different types of cell lines by changing the device dimensions.

We foresee interest in using this and comparable devices in systems biology applications, where it is important to measure cell responses upon dynamic drug stimulation and derive/distinguish mathematical models of the systems,^25^ and in synthetic biology, both to quantitatively characterize responses to inputs in synthetic gene networks^26^ and to implement external feedback control using microfluidics/microscopy platforms^15,27–29^.

## Author Contributions

AM: experimental work, data analysis, and manuscript writing. EP: experimental work and manuscript writing. ALR: experimental work and manuscript writing. AE: methodology. LM: methodology, supervision, manuscript writing and reviewing, funding.

## Acknowledgements

We thank Dr Mark Jepson and Alan Leard (Wolfson Imaging Facility, University of Bristol) for their support. This work was funded by the Engineering and Physical Sciences Research Council (grants EP/R041695/1 and EP/S01876X/1 to L.M.), by EC funding H2020 (FET OPEN 766840-COSY-BIO) to L.M., and by BrisSynBio, a BBSRC/EPSRC Synthetic Biology Research Centre (BB/L01386X/1) to L.M.

## References

1. Stefania Torino, Brunella Corrado MI and GC. PDMS-Based Microfluidic Devices for Cell Culture. Inventions. Published online 2018:1–14. doi:10.3390/inventions3030065

2. Bergmann S, Steinert M. From Single Cells to Engineered and Explanted Tissues: New Perspectives in Bacterial Infection Biology. Vol 319. Elsevier Ltd; 2015. doi:10.1016/bs.ircmb.2015.06.003

3. Ruppen J, Cortes-Dericks L, Marconi E, et al. A microfluidic platform for chemoresistive testing of multicellular pleural cancer spheroids. Lab Chip. 2014;14(6):1198–1205. doi:10.1039/c3lc51093j

4. Mehling M, Tay S. Microfluidic cell culture. Curr Opin Biotechnol. 2014;25:95–102. doi:10.1016/j.copbio.2013.10.005

5. Halldorsson S, Lucumi E, Gómez-sjöberg R, Fleming RMT. Biosensors and Bioelectronics Advantages and challenges of micro fl uidic cell culture in polydimethylsiloxane devices. Biosens Bioelectron. 2015;63:218–231. doi:10.1016/j.bios.2014.07.029

6. Craighead H, Yang P. From microfluidic application to nanofluidic phenomena issue Reviewing the latest advances in microfluidic and nanofluidic microenvironments w. Chem Soc Rev. 2018;(3). doi:10.1039/b909900j

7. Rothbauer M, Zirath H, Ertl P. Recent advances in microfluidic technologies for cell-to-cell interaction studies. Lab Chip. 2018;18(2):249–270. doi:10.1039/c7lc00815e

8. Klein AM, Mazutis L, Weitz DA, et al. Droplet Barcoding for Single-Cell Transcriptomics Applied to Embryonic Stem Cells Resource Droplet Barcoding for Single-Cell Transcriptomics Applied to Embryonic Stem Cells. Cell. 2015;161(5):1187–1201. doi:10.1016/j.cell.2015.04.044

9. Evan Z. Macosko, Anindita Basu, Rahul Satija, James Nemesh K, Shekhar, Melissa Goldman, Itay Tirosh, Allison R. Bialas, Nolan Kamitaki1 EM, Martersteck, John J. Trombetta, David A. Weitz5, Joshua R. Sanes9 AK, Shalek, Aviv Regev and SAM. Highly parallel genome-wide expression profiling of individual cells using nanoliter droplets. Cell. 2017;171(6):1437–1452. doi:10.1016/j.cell.2015.05.002.Highly

10. Mustafa A, Pedone E, Marucci L, Moschou D, Lorenzo M Di. A flow-through microfluidic chip for continuous dielectrophoretic separation of viable and non-viable human T-cells. Electrophoresis. Published online 2021:1–8. doi:10.1002/elps.202100031

11. Schuster B, Junkin M, Kashaf SS, et al. Automated microfluidic platform for dynamic and combinatorial drug screening of tumor organoids. Nat Commun. 2020;11(1):1–12. doi:10.1038/s41467-020-19058-4

12. Kolnik M, Tsimring LS, Hasty J. Vacuum-assisted cell loading enables shear-free mammalian microfluidic culture. Lab Chip. 2012;12(22):4732–4737. doi:10.1039/c2lc40569e

13. Nocera GM, Viscido G, Brillante S, Carrella S, Bernardo D. The VersaLive platform enables pipette-compatible microfluidic mammalian cell culture for versatile applications. bioRxiv. Published online 2021:1–22.

14. Yazdian Kashani S, Keshavarz Moraveji M, Bonakdar S. Computational and experimental studies of a cell-imprinted-based integrated microfluidic device for biomedical applications. Sci Rep. 2021;11(1):1–17. doi:10.1038/s41598-021-91616-2

15. Postiglione L, Napolitano S, Pedone E, et al. Regulation of Gene Expression and Signaling Pathway Activity in Mammalian Cells by Automated Micro fl uidics Feedback Control. ACS Synth Biol. Published online 2018. doi:10.1021/acssynbio.8b00235

16. Chew YT, Xia HM, Shu C. Fluid Micromixing Technology and Its Applications for Biological and Chemical Processes. In: Ibrahim F, Osman NAA, Usman J, Kadri NA, eds. 3rd Kuala Lumpur International Conference on Biomedical Engineering 2006. Springer Berlin Heidelberg; 2007:16–20.

17. Ward, Kevin and Fan ZH. Mixing in microfluidic devices and enhancement methods. J Micromech Microeng. 2015;25(9):1–33. doi:10.1088/0960-1317/25/9/094001.Mixing

18. Dettinger P, Frank T, Etzrodt M, et al. Automated Micro fl uidic System for Dynamic Stimulation and Tracking of Single Cells. Anal Chem. Published online 2018. doi:10.1021/acs.analchem.8b00312

19. Luo C, Zhu X, Yu T, et al. A fast cell loading and high-throughput microfluidic system for long-term cell culture in zero-flow environments. Biotechnol Bioeng. 2008;101(1): 190–195. doi:10.1002/bit.21877

20. Wang L, Ni XF, Luo CX, Zhang ZL, Pang DW, Chen Y. Self-loading and cell culture in one layer microfluidic devices. Biomed Microdevices. 2009;11(3):679–684. doi:10.1007/s10544-008-9278-0

21. Wray J, Kalkan T, Gomez-Lopez S, et al. Inhibition of glycogen synthase kinase-3 alleviates Tcf3 repression of the pluripotency network and increases embryonic stem cell resistance to differentiation. Nat Cell Biol. 2011;13(7):838–845. doi:10.1038/ncb2267

22. Parchem RJ, Ye J, Judson RL, et al. reprogramming. Cell Stem Cell. 2015;14(5):617–631. doi:10.1016/j.stem.2014.01.021.Two

23. An D, Kim K, Kim J. Microfluidic system based high throughput drug screening system for curcumin/TRAIL combinational chemotherapy in human prostate cancer PC3 cells. Biomol Ther. 2014;22(4):355–362. doi:10.4062/biomolther.2014.078

24. Ruzycka M, Cimpan MR, Rios-Mondragon I, Grudzinski IP. Microfluidics for studying metastatic patterns of lung cancer. J Nanobiotechnology. 2019;17(1):1–30. doi:10.1186/s12951-019-0492-0

25. Marucci L. Nanog dynamics in mouse embryonic stem cells: Results from systems biology approaches. Stem Cells Int. 2017;2017. doi:10.1155/2017/7160419

26. Marucci L, Santini S, di Bernardo M, di Bernardo D. Derivation, identification and validation of a computational model of a novel synthetic regulatory network in yeast. J Math Biol. 2011;62(5):685–706. doi:10.1007/s00285-010-0350-z

27. Pedone E, Postiglione L, Aulicino F, et al. A tunable dual-input system for on-demand dynamic gene expression regulation. Nat Commun. 2019;10(1):1–13. doi:10.1038/s41467-019-12329-9

28. Pedone E, De Cesare I, Zamora-Chimal CG, et al. Cheetah: A Computational Toolkit for Cybergenetic Control. ACS Synth Biol. 2021;10(5):979–989. doi:10.1021/acssynbio.0c00463

29. Khazim M, Postiglione L, Pedone E, Rocca DL, Zahra C, Marucci L. Towards automated control of embryonic stem cell pluripotency. IFAC-PapersOnLine. 2019;52(26):82–87. doi:10.1016/j.ifacol.2019.12.240

