## Supplementary information for "A novel single layer microfluidic device for dynamic stimulation, culture and imaging of mammalian cells"

### S1: FEM Analysis

Numerical simulations based on finite element methods are routinely used to aid the development of microfluidic devices by verifying the design and operating parameters. The governing equations for our model are given as.

$$\rho\frac{\partial u}{\partial t} + \rho(\boldsymbol{u_{fluid}}.\nabla)\,\boldsymbol{u_{fluid}} = \nabla\,.\,[-\mathbf{pI} + \mathrm{K}] + \boldsymbol{F} \qquad \textbf{(1)}$$

$$\rho\nabla.\,\boldsymbol{u_{fluid}} = \mathbf{0} \qquad \textbf{(2)}$$

$$\nabla.\,\boldsymbol{j_i} + \boldsymbol{u}.\nabla\mathbf{c}_i = \boldsymbol{R_i} \qquad \textbf{(3)}$$

$$\boldsymbol{j_i} = -\boldsymbol{D_i}\nabla_i \qquad \textbf{(4)}$$

Time-dependent simulations were setup to mimic the experiments. Flow velocities in the main channel and cell traps were calculated using the General Laminar Flow. The magnitude of flow velocity and shear rate in serpentines, flow channels and cell culture chambers are depicted in Figure S1. The flow velocity magnitude and the shear rate have their maximum values in the serpentines and minimum in the cell culture chambers. This ensures the claim that cells trapped in cell culture chambers are not exposed to excessive fluid shear stress enabling them to proliferate.

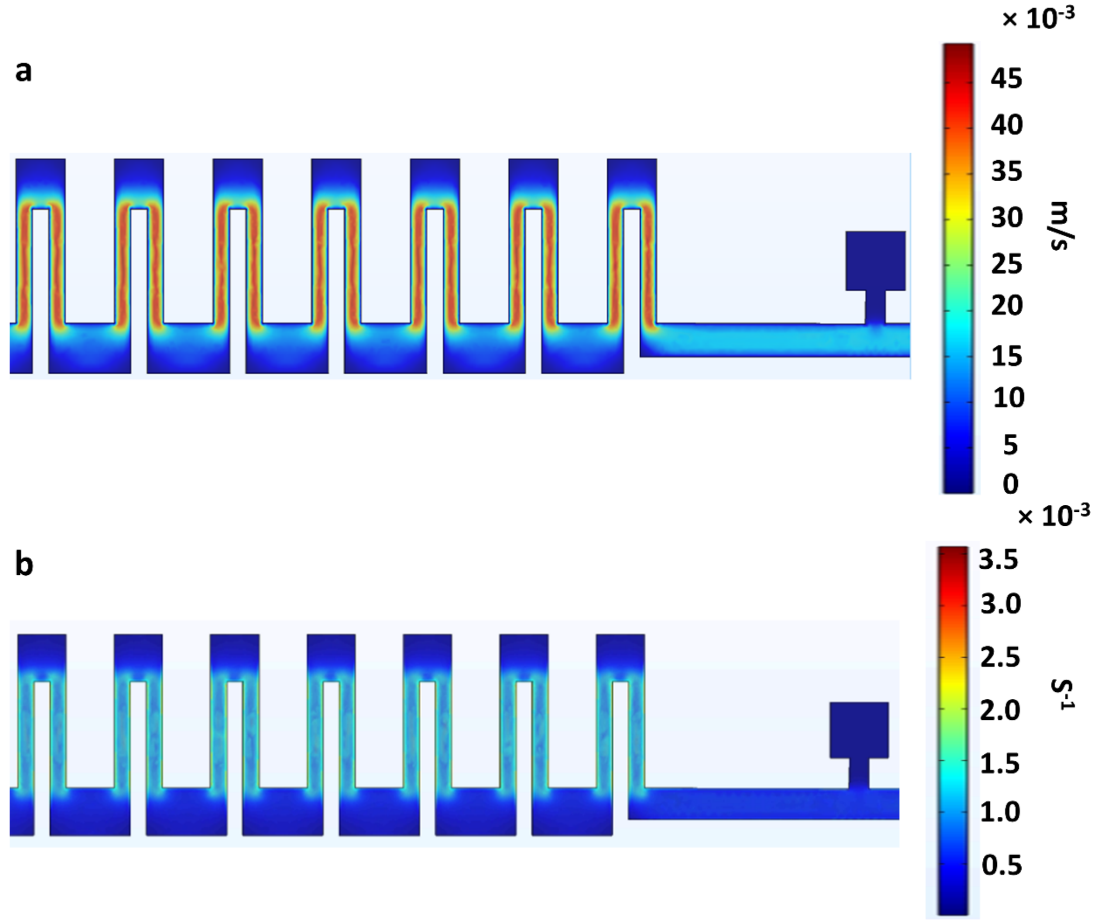


**Figure S1**: (a) Flow velocity (in m/s) in serpentine, flow channel and cell trapping chambers. (b) Shear rate (in $s^{-1}$) in serpentine, flow channel and cell trapping chambers.

### S2: Media exchange experiments

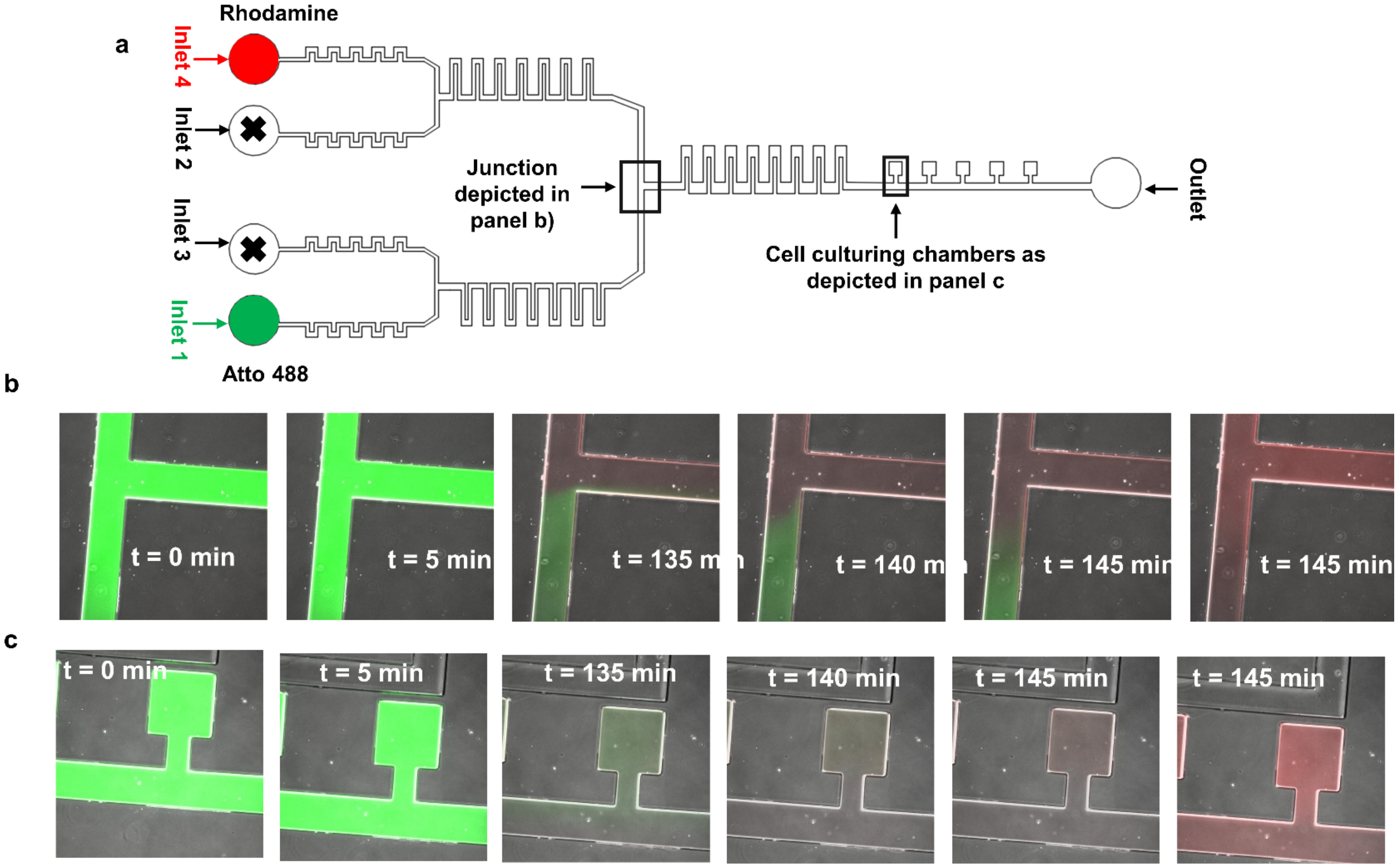


**Figure S2**: Media exchange experiments for extended time at 50 μLhr$^{-1}$. (a) Microfluidic device design with highlighted parts. (b) Images of the junction indicated in (a). At the start of the experiment, we have only green dye and at t = 135 min the inlet with green dye is switched off. The red dye starts to take over the junction and t=145 minutes it completely takes over the junction. (c) Images of the first cell culture chamber. At t=145 minutes it completely takes over the cell culture chamber.
